# Natural selection of glycoprotein B (gB) mutations that rescue the small plaque phenotype of a fusion-impaired herpes simplex virus (HSV) mutant

**DOI:** 10.1101/412221

**Authors:** Qing Fan, Sarah J. Kopp, Nina C. Byskosh, Sarah A. Connolly, Richard Longnecker

## Abstract

Glycoprotein B (gB) is a conserved viral fusion protein that is required for herpesvirus entry. To mediate fusion with the cellular membrane, gB refolds from a prefusion to a postfusion conformation. We hypothesize that an interaction between the C-terminal arm and the central coiled-coil of the herpes simplex virus 1 (HSV-1) gB ectodomain is critical for fusion. We previously reported that three mutations in the C-terminal arm (I671A/H681A/F683A, called gB3A) greatly reduced cell-cell fusion and that virus carrying these mutations had a small plaque phenotype and delayed entry into cells. By serially passaging gB3A virus, we selected three revertant viruses with larger plaques. These revertant viruses acquired mutations in gB that restore the fusion function of gB3A, including gB-A683V, gB-S383F/G645R/V705I/A855V, and gB-T509M/N709H. V705I and N709H are novel mutations that map to the portion of domain V that enters domain I in the postfusion structure. S383F, G645R, and T509M are novel mutations that map to an intersection of three domains in a prefusion model of gB. We introduced these second-site mutations individually and in combination into wild-type gB and gB3A to examine the impact of the mutations on fusion and expression. V705I and A855V (a known hyperfusogenic mutation) restored the fusion function of gB3A, whereas S383F and G645R dampened fusion and T509M and N709H worked in concert to restore gB3A fusion. The results identify two regions in the gB ectodomain that modulate the fusion activity of gB, potentially by impacting intramolecular interactions and stability of the prefusion and/or postfusion gB trimer.

**IMPORTANCE:** Glycoprotein B (gB) is an essential viral protein that is conserved in all herpesviruses and is required for virus entry. gB is thought to undergo a conformational change that provides the energy to fuse the viral and cellular membranes, however the details of this conformational change and the structure of the prefusion and intermediate conformations of gB are not known. Previously, we demonstrated that mutations in the gB “arm” region inhibit fusion and impart a small plaque phenotype. Using serial passage of a virus carrying these mutations, we identified revertants with restored plaque size. The revertant viruses acquired novel mutations in gB that restored fusion function and mapped to two sites in the gB ectodomain. This work supports our hypothesis that an interaction between the gB arm and the core of gB is critical for gB refolding and provides details about the function of gB in herpesvirus mediated fusion and subsequent virus entry.

## INTRODUCTION

Herpes simplex virus (HSV) causes recurrent mucocutaneous lesions on the mouth, face or genitalia and spread to the central nervous system which can lead to meningitis or encephalitis (1). Infection of host cells occurs by fusion of the virion envelope with a cell membrane to deliver the nucleocapsid and the viral genome into the host cell. HSV entry into cells and virus-induced cell-cell fusion require the coordinated action of the four viral entry glycoproteins: glycoprotein D (gD), gH, gL, and gB. The binding of gD to receptor results in a conformational change in gD that is proposed to signal the gHgL heterodimer to trigger the fusogenic activity of gB (2-4).

gB is a trimeric class III viral fusion protein that is conserved across all herpesviruses (5-7). Upon triggering, gB inserts into the cellular membrane and refolds from a prefusion to a postfusion conformation to bring the viral and cell membranes together. The structures of the postfusion forms of gB from three herpesviruses have been solved (5, 8-10). To date, attempts to capture a stable prefusion form of HSV-1 gB for crystallization have been unsuccessful (11). Alternative gB conformations have been modeled recently using cryoelectron tomography of gB present on membranes (12, 13). A prefusion model of gB has been developed using the crystal structure of the class III fusion protein from vesicular stomatitis virus (VSV) (14-16).

Our previous site-directed mutagenesis study demonstrated that fusion was impaired by mutations in an extended arm at the C-terminus of the ectodomain (domain V) that packs against the coiled-coil core of gB (domain III) in the post-fusion conformation (17). Alanine substitutions in three arm residues (I671A, H681A, and F683A; termed gB3A in this paper) that were predicted to disrupt interactions between the arm and coil greatly reduced fusion in a quantitative cell-cell fusion assay, indicating that the gB arm is important for fusion. Fusion function in gB3A was restored by the addition of a known hyperfusogenic mutation in the gB cytoplasmic tail, suggesting that the gB3A mutations did not cause global misfolding of gB. Due to the similarity of this gB coil-arm complex to the six-helix bundle of class I fusion proteins (18), we hypothesized that the gB coil-arm interactions were important for the transition from a prefusion to a postfusion gB conformation.

To further investigate the role of the gB arm region in virus entry, we generated and characterized HSV-1 carrying the gB3A mutations. This gB3A virus had small plaques, impaired growth, and delayed penetration into cells that was restored partially at elevated temperatures (19). These results supported our hypothesis that the gB3A mutations alter fusion kinetics by stabilizing a prefusion or intermediate conformation of gB or destabilizing the postfusion conformation.

In the present study, we passaged gB3A virus and selected for large plaque variants to identify mutations within gB that could restore gB3A fusion function. We selected three independent revertant viruses and sequencing revealed that all three viruses acquired mutations in gB. We cloned gB from the revertant viruses and introduced the mutations into wild-type (WT) gB and gB3A to analyze the effect of these mutations on cell surface expression and cell-cell fusion activity when coexpressed with WT gD, gH, and gL. The second-site revertant mutations identified two regions that modulate the fusion activity of gB3A: the C-terminus of the gB ectodomain and the intersection of three domains in a model of prefusion gB.

## RESULTS

### Selection of gB3A revertant viruses

Since gB3A virus exhibited delayed entry and a small plaque phenotype (19), we used serial passage to select for second-site mutations that would rescue the gB3A plaque morphology. We hypothesized that the location of these second-site mutations would reveal gB sites of functional importance, potentially including residues that affect the stability of gB and/or interact with the domain V arm region in a prefusion or intermediate gB conformation. Since our previous study showed that truncating the cytoplasmic tail of gB3A restored its fusion activity (17), we had reason to expect that second site mutations in gB would be able rescue the gB3A fusion defect.

Three independently purified stocks of DNA from a BAC carrying gB3A (pQF297) (19) were transfected individually into Vero cells expressing Cre recombinase to excise the BAC backbone. Virus harvested from these cells was amplified in Vero cells to generate three independent gB3A virus stocks. To select for revertant viruses possessing a growth advantage linked to a restoration of fusion function, these gB3A stocks were passaged serially in Vero cells. Virus harvested from each passage was titered to facilitate calculation of a multiplicity of infection (MOI) of 0.01 for the next passage. After serial passage, larger plaques were observed during titration and the infections spread faster in culture than the parental gB3A virus. For example, prior to passage 15, the gB3A virus stock designated 58621 required 7-15 days between passages to reach full cytopathic effect (CPE). By passage 20, CPE developed at the same rate for the 58621 infection and WT infections. By passage 24, the majority of plaques from the 58621 infection were large.

### Large plaque revertant viruses carry mutations in gB

A single large plaque revertant was selected from each of the three independently passaged gB3A stocks and the revertant virus was plaque purified. Viral DNA from the revertant viruses was used to clone and sequence the gB gene to allow us to examine the effect of gB mutations on fusion function in the absence of other viral genes that may have acquired mutations during the serial passaging. Sequencing results revealed that each revertant virus acquired distinct mutations within gB, including a single mutation A683V (isolated at passage 6), a quadruple mutation S383F/G645R/V705I/A855V (isolated at passage 25), and a double mutation T509M/N709H (isolated at passage 30).

### Fusion restoration in the gB3A-A683V revertant

The gB3A-A683V revertant virus demonstrated a larger plaque size than the parental gB3A virus, with a plaque morphology that resembled WT HSV (Fig. 1). The A683V mutation directly changes one of the original residues mutated in gB3A. Residue 683 is an alanine in gB3A and a phenylalanine in WT gB.

**Fig. 1.**
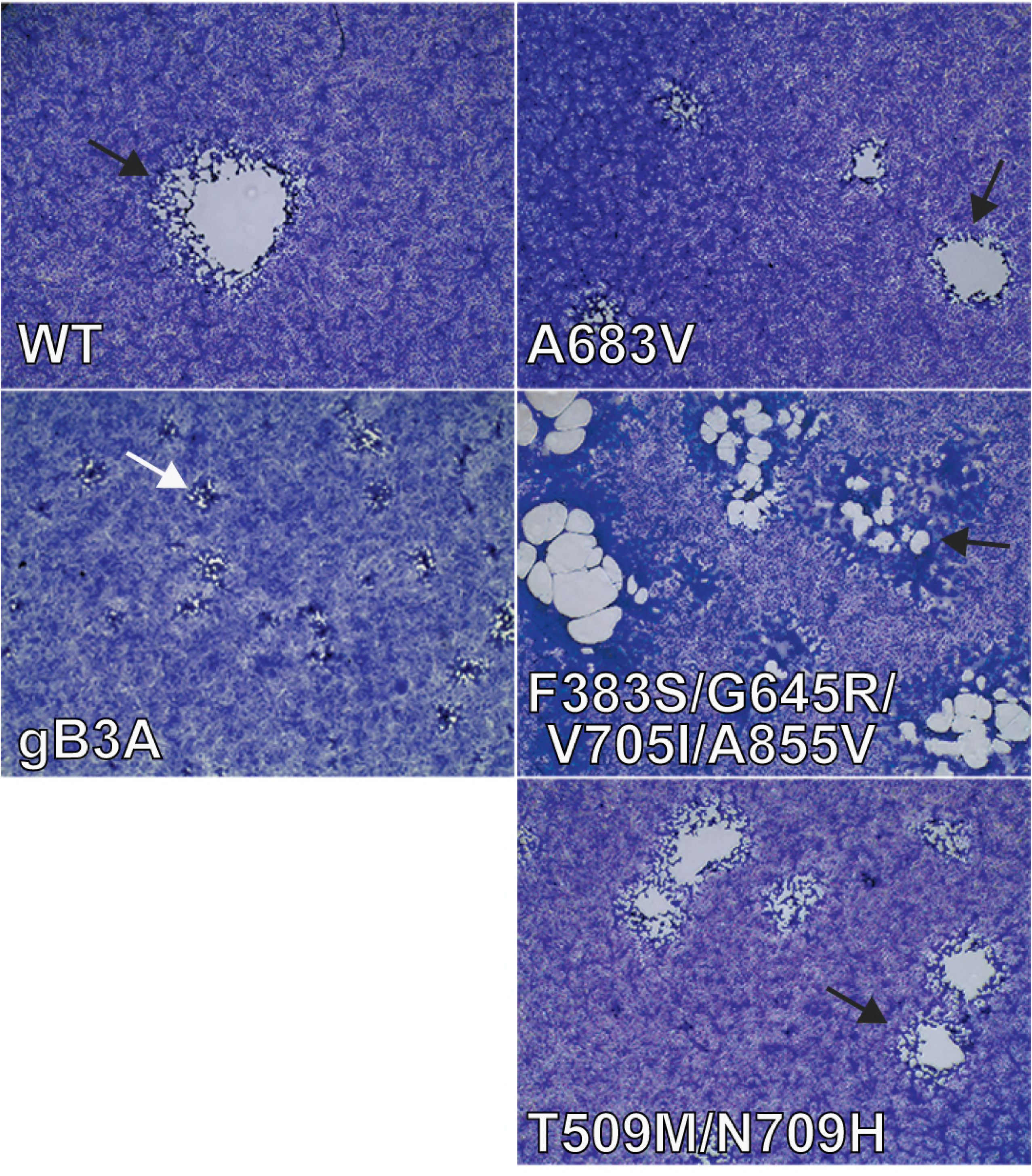
Plaque size and morphology of gB3A-revertant viruses. Vero cells were infected with BAC-derived WT HSV (GS3217), gB3A virus, or gB3A-revertant viruses (A683V, F383S/G645R/V705I/A855V, or T509M/N709H) at an MOI of 0.01. Three days post-infection, cells were stained and imaged at 4X magnification. gB3A plaques (white arrow) are compared to those of WT and revertant viruses (black arrows).

To assess the impact of the A683V mutation on fusion, the gB gene from this revertant virus was cloned into an expression vector (gB3A-A683V) with an N-terminal FLAG tag added to facilitate comparison of expression levels. This valine substitution at 683 also was added to a FLAG-tagged version of WT gB (gB-F683V). A virus-free cell-cell fusion assay was used to examine mutant gB fusion function. In this assay, one set of cells expressing T7 polymerase, gD, gH, gL, and version of gB is cocultured with a second set of cells expressing with an HSV receptor and luciferase under the control of the T7 promoter. Luciferase expression is used to quantify cell-cell fusion. As we previously reported, gB3A mediated nearly undetectable fusion (Fig. 2) (17). gB3A-A683V partially restored fusion, compared to gB3A, consistent with the restoration of plaque size exhibited by the revertant virus. gB-F683V showed reduced fusion compared to WT gB, indicating that, although a valine is functional in this position, a phenylalanine at this position promotes greater fusion. Interestingly, FLAG-tagged gB3A showed impaired cell surface expression compared to FLAG-tagged WT gB (Fig. 2). This effect on gB3A expression was unexpected because the FLAG-tagged WT gB and untagged WT gB have similar levels of surface expression (data not shown). The addition of the A683V mutation to FLAG-tagged gB3A restored expression, which may partially account for restoration of fusion function.

**Fig. 2.**
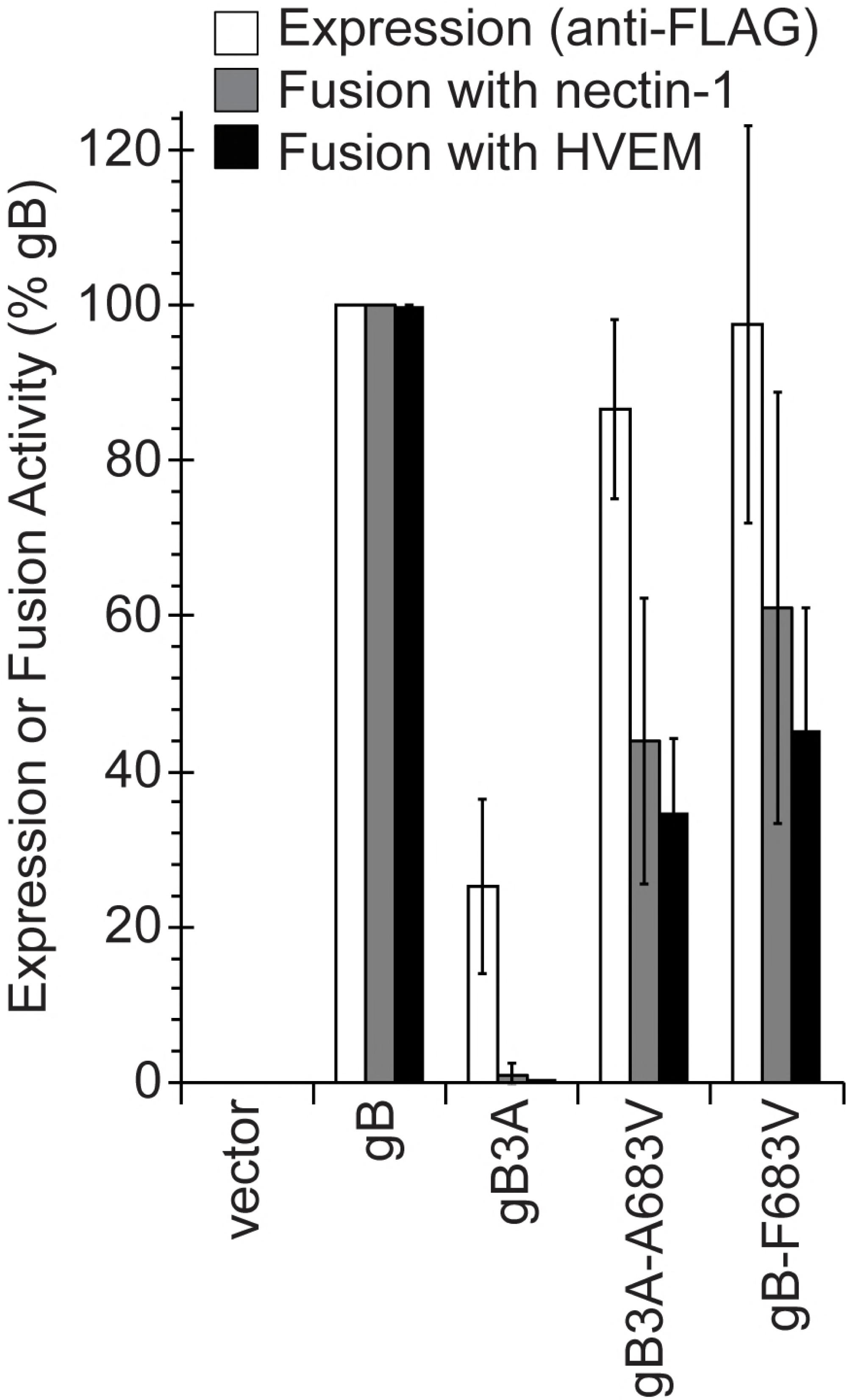
Expression of and fusion mediated by gB mutants constructs. gB residue 683 was mutated to valine in FLAG-tagged constructs of WT gB and gB3A. CHO-K1 cells (effector cells) were transfected with plasmids encoding T7 polymerase, gD, gH, and gL, plus either a version of gB or empty vector. Another set of CHO-K1 cells (target cells) was transfected with a plasmids encoding the luciferase gene under the control of the T7 promoter and either nectin-1 or HVEM receptor. One set of effector cells was used to determine cell surface expression of gB by CELISA using a MAb specific for the FLAG tag (white bars). Two duplicate sets of effector cells were co-cultured with the target cells expressing nectin-1 (gray bars) or HVEM (black bars) and luciferase activity was assayed as measure of cell-cell fusion activity. The results are expressed as a percentage of wild-type FLAG-gB expression or fusion activity, after subtracting background values measured in the absence gB expression (vector). The means and standard deviations of at least three independent experiments are shown.

### Fusion restoration in the gB3A-S383F/G645R/V705I/A855V revertant

The gB3A-S383F/G645R/V705I/A855V revertant virus demonstrated a larger plaque size than the parental gB3A virus, with a plaque morphology that resembles a syncytial virus (Fig. 1). This morphology agrees with previous work that identified gB-A855V as a syncytial mutation (20, 21).

The gB gene from this quadruple mutant was cloned into the same expression vector as above (gB3A-S383F/G645R/V705I/A855V) to assay the effect of these mutations on cell-cell fusion and cell surface expression. The revertant gB carrying the four second-site mutations exhibited nearly WT levels of both cell surface expression and fusion (Fig. 3A). This restoration of fusion is consistent with restored plaque size of the revertant virus. As with the A683V mutant, the effect of the second-site mutations on the surface expression of FLAG-gB3A may contribute to enhanced fusion.

**Fig. 3.**
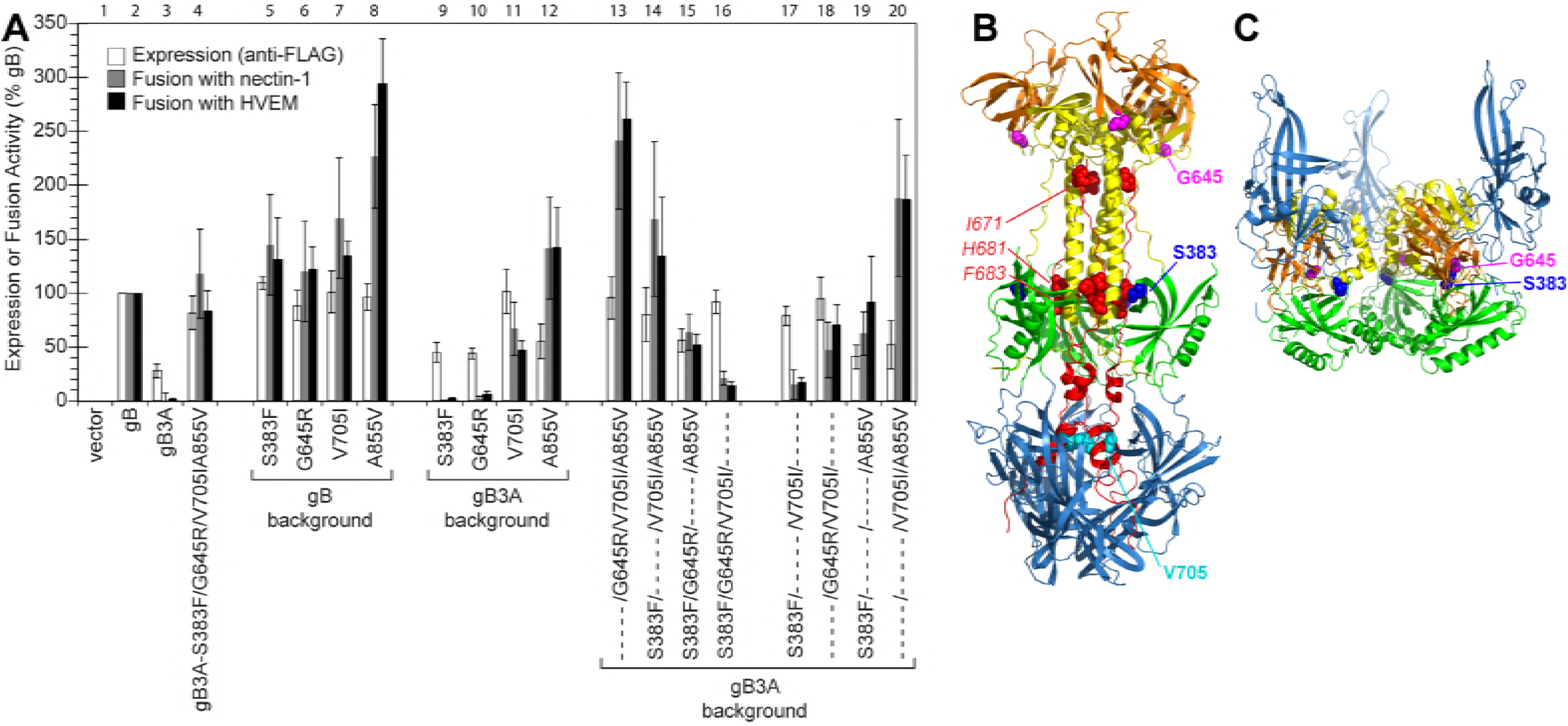
Expression of and fusion mediated by gB mutant constructs derived from gB3A-S383F/G645R/V705I/A855V. (A) gB mutations were added to FLAG-tagged WT gB or gB3A constructs, as indicated. Cell surface expression (white bars) and fusion activity (gray and black bars) of the gB constructs were assayed as in Fig. 2. Column sets are numbers above the graph. (B) Location of the mutated residues on postfusion gB. The crystal structure of the ectodomain of postfusion is gB (PDB ID 2GUM) is shown, with five domains colored, including DI (blue), DII (green), DIII (yellow), DIV (orange), and DV (red). The three residues mutated in DV of gB3A are shown (red spheres). Three of the mutations selected in the gB3A revertant are shown, including S383 (dark blue spheres), G645 (magenta spheres), and V705 (cyan spheres). A855 is not shown because it is located in the gB cytoplasmic tail. (C) Location of mutated residues on a model of prefusion gB. A model of the prefusion gB ectodomain is shown (14), with four domains colored as in part B. DV and the cytoplasmic tail are not present in the model, thus the residues mutated in DV of gB3A, V705, and A855 are not shown.

Sequencing the gB gene from earlier passages of this revertant virus that were not plaque purified showed that the G645R and A855V mutations (present at passage 8) appeared before the S383F mutation (present at passage 10) and that the V705I mutation appeared last (present in the final passage 25).

To examine how the four amino acid changes in the revertant virus may contribute to the restoration of plaque size, the mutations were introduced into gB singly or in combination. The four mutations were introduced individually into FLAG-tagged WT gB or gB3A. Then the four second-site mutations present in the revertant gB construct were mutated individually back to their corresponding WT residues. Four additional double mutations were also generated in the gB3A background.

Introduction of any one of the second-site mutations into WT gB did not affect the cell surface expression levels (Fig. 3A). As expected, A855V enhanced cell-cell fusion when added to WT gB (Fig. 3A, column 8). The A855V mutation was shown previously to enhance cell-cell fusion (22, 23). Individual introduction of the other three second-site mutations into WT gB enhanced fusion only modestly.

The addition of S383F or G645R to gB3A did not alter cell surface expression or fusion substantially. The addition of A855V to gB3A enhanced fusion to greater than WT levels (Fig. 3A, column 12), similar to the results observed in the WT gB background. Interestingly, V705I enhanced both gB3A cell surface expression and fusion (Fig. 3A, column 11).

When the second-site mutations present in gB3A-S383F/G645R/V705I/A855V were restored individually to the corresponding WT gB residues, loss of the A855V mutation resulted in greatly reduced fusion levels (Fig. 3A, column 16), as expected. Loss of the V705I mutation modestly reduced both expression and fusion (column 15). Loss of either the S383F or G645R second-site mutations (columns 13-14) did not impact expression levels, but surprisingly fusion was enhanced compared to the revertant protein carrying all four mutations (column 4). This finding suggests that S383F or G645R independently dampen fusion mediated by this revertant gB.

To further examine the role of these second-site mutations, four gB3A constructs carrying double mutations were created (Fig. 3A). The gB3A construct carrying both the V705I and A855V mutations showed the highest levels of fusion (Fig. 3A, column 20), higher than those observed when A855V was added to gB3A alone (column 12), suggesting that V705I promotes fusion in addition to A855V. A comparison of gB3A-V705I/A855V (column 20) with the revertant carrying all four mutations (column 4) confirms that S383F and G645R dampen fusion in this revertant. Similarly, a dampening effect of S383F on fusion is apparent when comparing gB3A-S383F/A855V (column 19) with gB3A-A855V (column 12) or gB3A-S383F/V705I (column 17) with gB3A-V705I (column 11).

### Fusion restoration in the gB3A-T509M/N709H revertant

The gB3A-T509M/N709H revertant virus demonstrated larger plaques than the parental gB3A virus, with a plaque morphology resembling WT HSV (Fig. 1). Both of these mutations are in the gB ectodomain and neither were previously reported to be hyperfusogenic. Interestingly, in the postfusion structure of gB, T509 is a contact residue for F683, located in the central trimeric coiled-coil. To examine the impact of these mutations on gB expression and fusion, the gB gene from this virus was cloned into the same expression vector as above (gB3A-T509M/N709H). The addition of these two mutations to gB3A enhanced fusion (Fig. 4), consistent with the larger plaque phenotype of this virus. As with the previous two revertant gB constructs, the expression of the revertant gB was enhanced compared to FLAG-gB3A, which may contribute to improved fusion function. When the mutations were added individually to WT gB, neither mutation enhanced fusion (Fig. 4), indicating that these mutations are not hyperfusogenic like A855V. In fact, both of the mutations somewhat decreased fusion mediated by WT gB. When the mutations were added individually to gB3A, no fusion was detected (Fig. 4), indicating that neither mutation alone can rescue the fusion defect imparted by gB3A. The mutations act in combination to restore fusion mediated by gB3A and are of particular interest because they have not been previously described and they map to two sites in the prefusion and postfusion gB structures that correspond with the gB3A-S383F/G645R/V705I/A855V revertant.

**Fig. 4.**
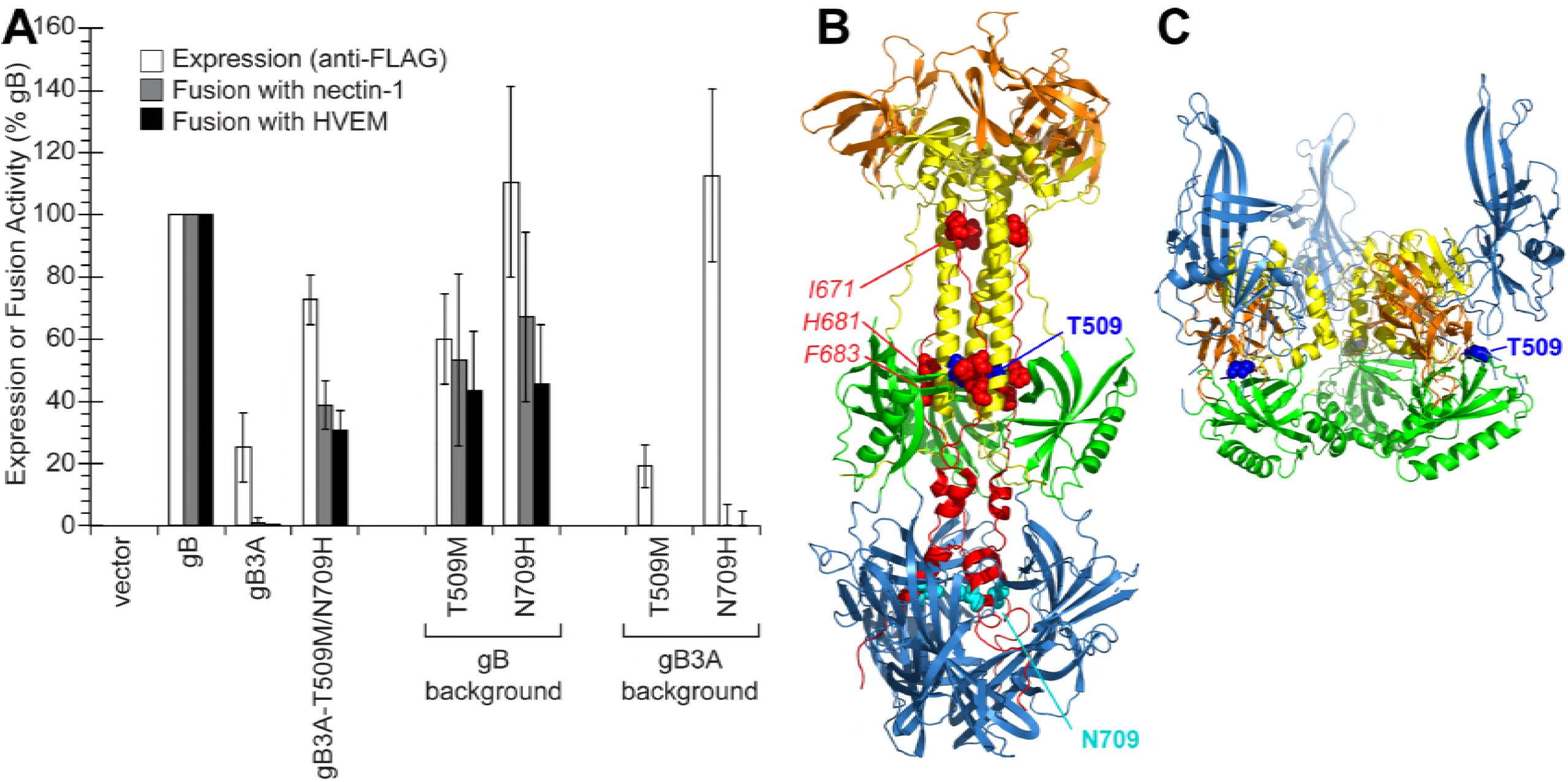
Expression of and fusion mediated by gB mutant constructs derived from gB3A-T509M/N709H. (A) gB mutations were added to FLAG-tagged WT gB or gB3A constructs, as indicated. Cell surface expression (white bars) and fusion activity (gray and black bars) of the gB constructs were assayed as in Fig. 2. (B) Location of mutated residues on postfusion gB. The structure of postfusion gB (PDB ID 2GUM) is shown, with the gB3A mutated residues (red spheres) and five domains colored as in Fig. 3B. The two mutations selected in the gB3A revertant are shown, including T509 (dark blue spheres) and N709 (cyan spheres). (C) Location of mutated residues on a model of prefusion gB. A model of prefusion gB is shown (14), with domains colored as in Fig. 2C. T509 (blue spheres) is shown. DV is not present in the model, thus N709 is not shown.

## DISCUSSION

The gB3A mutations (I671A/H681A/F683A) were designed based on the postfusion structure of gB (5) to reduce interactions between the domain V arm and the central coiled-coil (17). We previously showed that these mutations inhibited cell-cell fusion and virus carrying these mutations exhibited a growth defect and formed 200-fold smaller plaques than WT virus (19). To investigate how these gB3A mutations inhibit fusion, we serially passaged BAC-derived gB3A virus to select for mutations that overcome the fusion defect and restore a larger plaque size. Three revertant viruses were isolated and all three viruses carried mutations in gB3A.

To examine whether the newly acquired mutations in gB3A could account for the restored plaque size, we cloned the gB genes from these viruses into expression vectors. The second-site mutations enhanced gB3A cell-cell fusion, suggesting that these mutations were responsible for the larger plaque size. Thus, natural selection identified several second-site gB mutations that restore gB3A function. Although additional mutations outside of gB also may have contributed to the restored plaque size and/or growth of the revertant viruses, cloning the revertant gB genes allowed us to analyze the effect of the gB mutations in isolation when coexpressed with WT gD, gH, and gL.

The gB3A-A683V virus acquired a change in residue 683, one of the residues originally mutated to alanine in gB3A. Selection of this revertant mutation underscores the importance of residue 683 to gB fusion function and demonstrates that, although in WT residue 683 is a phenylalanine, a valine at this position is sufficient to promote fusion better than alanine, an amino acid with a smaller side-chain (Fig. 2A).

This A683V mutation was selected independently twice upon serial passage of gB3A virus (data not shown). Interestingly, among the three original mutation sites, A683 was the only one changed. In postfusion WT gB, I671, H681, and F683 pack against the central coiled-coil of gB (Fig. 3B). The selection pressure for a substitution at gB3A A683 rather than A681 can be explained by the structure, but the reason an A683V mutation was selected over a revertant substitution in I671 is unclear. I671 and F683 contact the coil more extensively than H681, with I671 and F683 contacting six and five coil residues, respectively, whereas H681 contacts only three coil residues (5). The side-chain atoms of I671 and F683 make 14 and 13 contacts with the coil, respectively, whereas the side-chain atoms of H681 make only seven contacts. The specific importance of I671 and F683 was apparent when we previously showed that individually mutating F683 or I671 alanine impairs fusion more than mutating H681 (17).

The gB3A-S383F/G645R/V705I/A855V virus retained the original three alanine mutations and acquired four additional mutations, including A855V, a known hyperfusogenic mutation located in the cytoplasmic tail of gB (22). By analyzing individual mutations, we confirmed that A855V alone enhances both gB3A and WT gB fusion (Fig. 3A). A855V and G645R mutations were the first mutations acquired in the revertant virus, present by passage 8. The larger plaque phenotype was apparent at this early passage, consistent with A855V being primarily responsible for the large plaque phenotype. The structure of the gB cytoplasmic tail was solved recently (24) and it forms a trimer that is proposed to stabilize prefusion gB, functioning as a clamp. Residue 855 lies within a long helix that angles upwards toward the membrane. Mutation of this residue may enhance fusion by weakening the trimerization of the cytoplasmic domain. Alternatively, this residue is on the periphery of the cytoplasmic domain and it could serve as a site of interaction with the gH cytoplasmic tail (24).

Interestingly, S383F or G645R independently appear to dampen fusion. This may seem counterintuitive initially, however maximal levels of fusion do not correlate necessarily with maximal titers. For example, virus complemented with hyperfusogenic gB yields lower viral titers than WT gB (17, 25, 26). The S383F and G645R mutations may provide a selective advantage to the revertant virus by preserving infectious virus production in the presence of the hyperfusogenic A855V mutation and the boosting titer of this virus. Although S383 and G645 are far apart in the postfusion structure of gB (Fig. 3B), they are located close to one another in a prefusion model of gB (Fig. 3C) (14), is based on the prefusion structure of the VSV fusion protein (15). S383 and G645 sit at the intersection of domains II, III, and IV, suggesting the possibility that mutations in these residues impact interdomain interactions that stabilize prefusion gB.

G645 lies in the middle of a stretch of three glycines. Recently, a mutation very similar to G645R was selected by serial passage of a gL-null pseudorabies virus (PRV) strain (27). gL-null PRV can spread cell-to-cell and passage of this virus selected for several second site mutations, including PRV gB G672R. This PRV gB mutation lies in a residue that corresponds to the HSV-1 gB glycine immediately adjacent to G645. PRV gB G672R was shown to be a modestly hyperfusogenic mutation when added to WT PRV gB.

In the postfusion gB structure, S383 maps in the same plane as residues H681 and F683 (Fig. 3B), presenting the possibility that replacing this small polar serine with a large hydrophobic phenylalanine may impact local interactions and alter how the arm packs against the coiled-coil in gB3A.

The V705I mutation enhanced surface expression levels, restoring the gB3A expression defect (Fig. 3A). Greater surface expression may account for some of the restoration of fusion function observed when V705I was added to gB3A. V705I also appears to boost to gB3A fusion function directly, since the addition of V705I/A855V to gB3A resulted in greater levels of fusion than the addition of A855V alone, despite similar levels of surface expression for both constructs (Fig. 3A, columns 12 and 20). V705I does not impact WT gB function, indicating that the addition of this larger hydrophobic side chain enhances gB3A fusion in a specific manner. Interestingly, the revertant virus acquired the V705I mutation after the other three mutations. The addition of V705I boosts cell-cell fusion activity compared to gB3A-S383F/G645R/A855V (Fig. 3A, columns 4 and 15), bringing it to the WT gB levels, potentially the optimal level for virus replication. V705 lies in domain V of gB, downstream from the three alanine mutations. In the postfusion structure of gB, V705 enters the core of domain I, the domain that includes the fusion loops and interacts with the host cell membrane (Fig. 3B). Mutation of V705 may influence the conformation of domain I or the refolding of gB3A into a postfusion conformation.

Like the quadruple mutant revertant virus, the gB3A-T509M/N709H revertant virus also retained the original three alanine mutations and acquired additional mutations that restored fusion function. Unlike the previous revertant, both T509M and N709H were required to restore gB3A function and neither mutation was hyperfusogenic in WT gB background. Remarkably, N709 is located near V705 in the postfusion structure (Fig. 4B), in the portion of domain V that penetrates domain I. The location of this mutation supports the importance of this region for gB fusion function. Similar to V705I, N709H enhanced gB3A expression specifically, suggesting a specific effect of this region on the gB3A structure.

T509 is a contact residue for F683 and is located in the central trimeric coiled-coil of postfusion gB (Fig. 4B). Although T509M did not restore gB3A fusion on its own, the substitution of a larger hydrophobic side chain at this position may partially restore an interaction between the gB arm and coil. Strikingly, in the prefusion gB model, T509 lies in at the intersection of domains II, III, and IV (Fig. 4C), near the location of S383 and G645 (Fig. 3C). Thus, mutations in this region were selected independently in two revertant viruses, suggesting that the interdomain interactions at this site may impact gB refolding during fusion.

We hypothesized that an interaction between the arm and coil regions of gB is important for gB refolding and we demonstrated that fusion is impeded by gB3A mutations that were designed to disrupt that interaction. Using *in vitro* evolution to select revertant viruses, we identified two gB regions that influence gB3A, including the C-terminal region of domain V in postfusion gB (residues I705 and N709) and the intersection of domains II, III, and IV in prefusion gB (residues S383, G645, and T509). Mutations at these residues may compensate for the gB3A fusion defect by destabilizing gB3A to reduce a kinetic barrier to fusion and/or enhancing gB expression. To investigate how other viral proteins regulate and/or trigger gB fusion activity, future work will select for revertant mutations outside of gB by passaging gB3A virus in cells that express gB3A as a cellular protein.

## MATERIALS AND METHODS

### Cells and antibodies

Chinese hamster ovary (CHO-K1; American Type Culture Collection (ATCC, USA)) cells were grown in Ham’s F12 medium supplemented with 10% fetal bovine serum (FBS) (ThermoFisher Scientific, USA). Vero cells (ATCC, USA) and Vero-cre cells that express Cre recombinase (kindly provided by Dr. Gregory Smith at Northwestern University) were grown in Dulbecco modified Eagle medium (DMEM) supplemented with 10% FBS, penicillin and streptomycin. The anti-FLAG MAb F1804 (Sigma) was used to assay cell surface expression.

### Plasmids and BACs

Previously described plasmids expressing HSV-1 KOS strain gB (pPEP98), gD (pPEP99), gH (pPEP100) and gL (pPEP101) (28), as well as nectin-1 (pBG38) (Geraghty et al, 1998) and HVEM (pBEC10) (29), were provided by Dr. Spear at Northwestern University. Plasmid pQF112 encodes an N-terminally FLAG-tagged version of WT HSV-1 gB (FLAG-gB) (30). Previously described BACs used in this study include a WT HSV-1 strain F BAC (GS3217; kindly provided by Dr. Gregory Smith, Northwestern University) and a BAC derived from GS3217 that encodes gB3A (pQF297) (19). Both BACs carry the red fluorescence protein (RFP) TdTomato under a cytomegalovirus promoter.

For this study, all gB mutants were cloned into the pFLAG-myc-CMV-21 expression vector (E5776; Sigma), substituting the gB signal sequence and adding an N-terminal FLAG epitope. A FLAG-tagged gB3A construct with gB3A mutations (I671A/H681A/F683A; pQF302) was subcloned from pSG5-HSVgB-I671A/H681A/F683A (17). Three revertant gB constructs were generated by amplifying the gB gene from viral DNA isolated using a DNeasy Blood and Tissue Kit (Qiagen, USA), including I671A/H681A/F683V (pQF338), I671A/H681A/F683A/S383F/G645R/V701I/A855V (pQF343), and I671A/H681A/F683A/T509M/N709H (pQF339). Quikchange site-directed mutagenesis (ThermoFisher Scientific, USA) was used to introduce specific mutations into FLAG-tagged gB3A (pQF302), including S383F (pQF319), G645R (pQF320), V705I (pQF321), A855V (pQF322), S383F/V705I (pQF328), G645R/V705I (pQF327), S383F/A855V (pQF324), V705I/A855V (pQF325). The mutant constructs G645R/V701I/A855V (pQF309), S383F/V705I/A855V (pQF310), S383F/G645R/A855V (pQF311), and S383F/G645R/V705I (pQF312) were created by using Quikchange on pQF343. Quikchange also was used to introduce mutations into FLAG-tagged WT gB (pQF112), including S383F (pQF305), G645R (pQF306), V705I (pQF307), A855V (pQF309), T509M (pQF367), and N709H (pQF369), and F683V (pQF368). The gB open reading frame was verified by sequencing for all clones.

### Virus stocks created from BAC DNA

WT HSV-1 BAC DNA (GS3217) and three independent stocks of gB3A virus BAC DNA (pQF297) were purified. The BAC DNA was transfected using Lipofectamine 2000 (Invitrogen, Carlsbad, CA) into Vero cells expressing Cre recombinase to excise the LoxP flanked BAC backbone, as previously described (19). The transfected cells were harvested and sonicated after one week for the WT BAC and three weeks for the gB3A BAC samples, based on when the majority of cells showed RFP expression. The harvested stocks were passaged once on Vero cells in roller bottles to generate virus stocks, including a WT stock and three independent gB3A stocks designated 58621, 57621, and 58632. For 58632, the virus stock was purified using three rounds of plaque purification using limiting dilution in 96 well plates.

### Selection of gB3A-revertant viruses

Vero cells were infected with the gB3A virus stocks at an MOI of 0.01. When full disruption of the monolayer by CPE was observed, cells were harvested and sonicated to create a virus stock. Virus then was reseeded on Vero cells at an MOI of 0.01 for serial passage. For the gB3A stocks 57621 and 58621, virus was passaged in 6 well plates for 18 passages and then passaged in roller bottles. For gB3A stock 58632, virus was passaged directly in roller bottles. Virus stocks were titered at each passage to facilitate calculation of an MOI of 0.01 for the next passage. When the titering step revealed large plaques, virus from a single large plaque was purified through three rounds of plaque purification using limiting dilution in 96-well plate. Prior to the emergence of larger plaques, each gB3A virus passage took 14-21 days. After larger plaques were observed, full CPE was achieved at 3-4 days post-infection, similar to a WT virus infection. The gB gene from larger plaque (revertant) virus stocks was sequenced. At passage 6, a 58632 revertant virus carrying the gB mutation A683V was isolated. At passage 25, a 58621 revertant virus carrying gB mutations S383F/G645R/V705I/A855V was isolated. At passage 30, a 57621 revertant virus carrying gB mutations T509M/N709H was isolated.

### Microscopy of plaque morphology

Plaques were visualized using Giemsa (Sigma-Aldrich, USA) staining after 3 days of infection and imaged with transmitted light microscopy using EVOS Cell Imaging Systems at 4x.

### Cell-cell fusion assay

The fusion assay was performed as previously described (28). Briefly, CHO-K1 cells were seeded in 6-well plates overnight. One set of cells (effector cells) were transfected with 400 ng each of plasmids encoding T7 RNA polymerase, gD, gH, gL, and either a gB construct or empty vector, using 5 μl of Lipofectamine 2000 (Invitrogen, USA). A second set of cells (target cells) was transfected with 400 ng of a plasmid encoding the firefly luciferase gene under control of the T7 promoter and 1.5 μg of either receptor (HVEM or nectin-1) or empty vector, using 5 μL of Lipofectamine 2000. After 6 h of transfection, the cells were detached with versene and resuspended in 1.5 mL of F12 medium supplemented with 10% FBS. Effector and target cells were mixed in a 1:1 ratio and re-plated in 96-well plates for 18 h. Luciferase activity was quantified using a luciferase reporter assay system (Promega) and a Wallac-Victor luminometer (Perkin Elmer).

### Cell-based ELISA (CELISA)

To evaluate the cell surface expression of gB mutants, CHO-K1 cells seeded in 96-well plates were transfected with 60 ng of empty vector or a gB construct using 0.15 μl of Lipofectamine 2000 (Invitrogen) diluted in Opti-MEM (Invitrogen). After 24 h, the cells were rinsed with phosphate buffered saline (PBS) and CELISA staining was performed as previously described (31), using the primary anti-FLAG MAb F1804. After incubation with primary antibody, the cells were washed, fixed, and incubated with biotinylated goat anti-mouse IgG (Sigma), followed by streptavidin-HRP (GE Healthcare) and HRP substrate (BioFX).

## ACKNOWLEDGMENTS

We thank Dr. Gregory Smith for providing us with the HSV-1 BAC GS3217 and Dr. Yasushi Kawaguchi for providing the parental BAC. We thank Nan Susmarski for timely and excellent technical assistance and members of the Longnecker laboratory for their help in these studies. Sequencing services were performed at the Northwestern University Genomics Core Facility. R.L. is the Dan and Bertha Spear Research Professor in Microbiology-Immunology. This work was supported by NIH grant CA021776 to R.L.

